# Flow-induced symmetry breaking in growing bacterial biofilms

**DOI:** 10.1101/627208

**Authors:** Philip Pearce, Boya Song, Dominic J. Skinner, Rachel Mok, Raimo Hartmann, Praveen K. Singh, Hannah Jeckel, Jeffrey S. Oishi, Knut Drescher, Jörn Dunkel

**Affiliations:** Department of Mathematics, Massachusetts Institute of Technology, Cambridge MA 02139-4307, USA; Department of Mechanical Engineering, Massachusetts Institute of Technology, Cambridge, MA 02139-4307, USA; Max Planck Institute for Terrestrial Microbiology, 35043 Marburg, Germany; Department of Physics, Bates College, Lewiston, ME 04240, USA; Department of Physics, Philipps-Universität Marburg, 35043 Marburg, Germany

## Abstract

Bacterial biofilms represent a major form of microbial life on Earth and serve as a model active nematic system, in which activity results from growth of the rod-shaped bacterial cells. In their natural environments, ranging from human organs to industrial pipelines, biofilms have evolved to grow robustly under significant fluid shear. Despite intense practical and theoretical interest, it is unclear how strong fluid flow alters the local and global architectures of biofilms. Here, we combine highly time-resolved single-cell live imaging with 3D multi-scale modeling to investigate the mechanisms by which flow affects the dynamics of all individual cells in growing biofilms. Our experiments and cell-based simulations reveal three quantitatively different growth phases in strong external flow, and the transitions between them. In the initial stages of biofilm development, flow induces a downstream gradient in cell orientation, causing asymmetrical droplet-like biofilm shapes. In the later developmental stages, when the majority of cells are sheltered from the flow by the surrounding extracellular matrix, buckling-induced cell verticalization in the biofilm core restores radially symmetric biofilm growth, in agreement with predictions of a 3D continuum model.

## INTRODUCTION

Fluid flow is a key element of many natural and industrial environments in which bacteria form biofilms, from rivers [1], pipes [2] and filtration devices [3] to the human heart [4], intestines [5] and mouth [6], where biofilms cause a major economic and health burden. Hydrodynamic effects have been found to play a crucial role during the initial attachment of cells to surfaces [7]. Later in development, flow provides nutrients to surface-attached biofilm communities, while removing metabolic waste products and signaling molecules [8–10]. There is therefore strong practical and theoretical interest in understanding the interaction between biofilms and external flow fields [8, 11]. Identifying the multi-scale dynamics of such growth-active nematics under the influence of shear will be helpful when adapting current theories for active matter [12, 13] to describe and predict bacterial biofilm growth across model systems and species [14, 15].

Imposed fluid shear has been observed to produce striking aerofoil-like shapes [16–18] during the early stages of biofilm growth and, in some cases, long filaments or streamers that extend far downstream [1, 19]. It has often been assumed that the key driver of the observed architecture of biofilms in flow is bulk deformation or erosion of biofilm biomass [16, 19, 20]. Recently, new imaging methodologies were developed to quantify biofilm dynamics at single-cell resolution, yet these studies have focused on conditions with very low flow [17, 21–23]. Earlier work that quantified biofilm architecture in high flow did not resolve the microscopic dynamics [1, 19], or did not explain the mechanisms by which high shear modifies the microcoscopic and macroscopic biofilm architecture [17]. Despite the new imaging techniques, and the extensive environmental relevance of flow-biofilm interactions, it has therefore remained unclear how flow reorients cells in space and time during biofilm growth, and, in turn, how these microscopic cellular reorientations contribute to the overall biofilm morphogenesis.

Here, we investigate comprehensively the effects of fluid shear on individual cell dynamics within growing *Vibrio cholerae* biofilms, by combining multi-scale modeling with highly time-resolved imaging at single-cell resolution (Fig. 1). First, we establish the translational and orientational dynamics of cells within early-stage biofilm microcolonies in flow, by constraining an individual cell-based model with the imaging data. Subsequently, these dynamics are included in a minimal continuum model that identifies the physical processes necessary to explain the biofilm architectural development observed at the later stages. We find that the bulk biofilm dynamics are determined almost entirely by cellular orientations inside the biofilm, representing the local flow-induced nematic director field, rather than by biofilm deformation or cell erosion as has been previously hypothesized [8, 17].

**FIG. 1.**
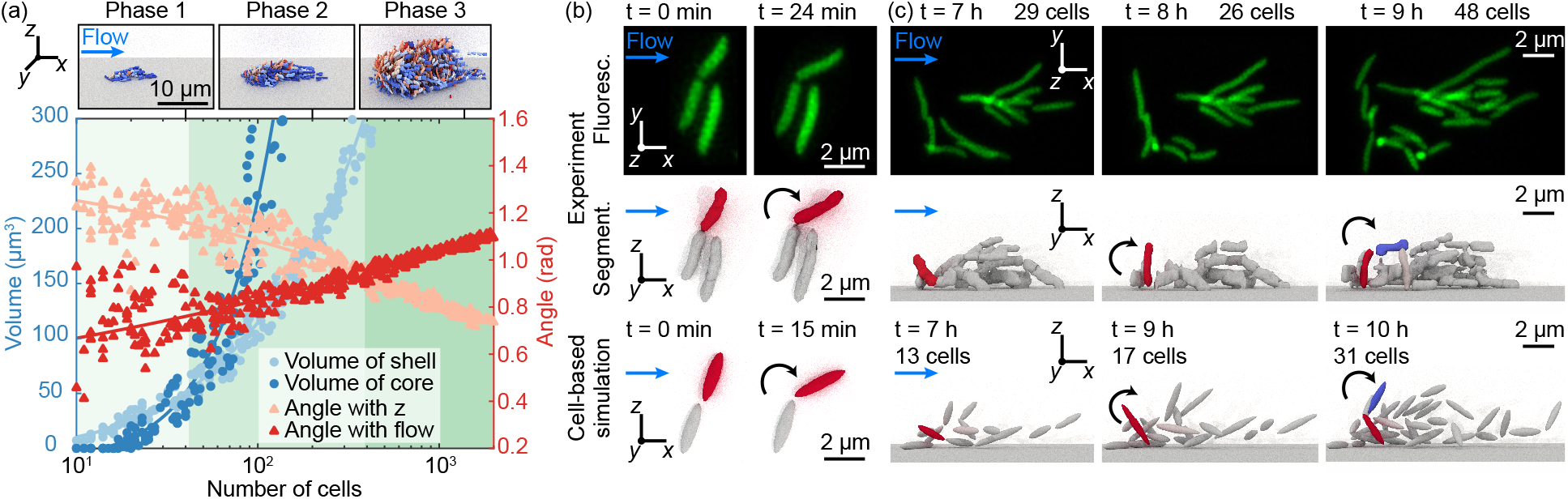
The three phases of *V. cholerae* biofilm growth and hydrodynamic cell alignment mechanisms in strong flow. (a) The transition from the 2D surface-dominated growth phase 1 (light green) to the 3D bulk-dominated phase 2 (green) occurs when the volume of the biofilm shell (light blue circles) equals the volume of the biofilm core (dark blue circles); see SM [24] for details of the volume measurements. The transition to the verticalization phase 3 (dark green) occurs when the average cell-orientation angles with *z* (light red triangles) and the flow direction (dark red triangles) cross. The diagram shows the combined data from *n* = 3 independent biofilm experiments, with typical snapshots of biofilms in each phase above (Movie 1). (b,c) Flow-induced cell re-orientation dynamics in the initial phase of biofilm growth. (b) Cells in direct contact with the surface align with the flow as a result of a torque generated by a combination of the fluid drag and asymmetrical attachment to their parent cell at the pole; see also Fig. S10 for additional intermediate snapshots. Fluorescence images show projection of a confocal *z*-stack. (c) Cells at the front of the biofilm (red) align vertically as a result of the torques *τ*_drag_ and *τ*_shear_; see also Fig. S11. When a vertically-oriented cell (red) divides, the daughter cell (blue), if exposed to shear, aligns with the flow.

## RESULTS

To investigate the effects of flow on biofilms in a broadly applicable setting, we imaged biofilm development on glass surfaces at cellular resolution in a flow channel with a shear rate of 2000 s^*−*1^ (Re ≈ 1), which is a typical order of magnitude for flows in natural and man-made environments containing bacteria [25–27].

To achieve the required time resolution, we improved a recently developed single-cell live-imaging technique, whereby adaptive confocal microscopy is combined with ground-truth-calibrated 3D image segmentation suitable for our model organism: *V. cholerae* with a straight cell shape (∆*crvA*), constitutively expressing a green fluorescent protein (sfGFP) (Fig. 1b,c; SM [24]). The increased time resolution, now continuously at ∆*t* = 6 min, enabled us to visualize previously inaccessible transient cell reorientations by the flow (Fig. 1b,c), in addition to reconstructing cell lineages and measuring cellular growth rates. By combining these new data with previously obtained data for larger biofilms [17], we obtained a comprehensive dataset showing the effects of fluid flow in unprecedented detail on biofilms growing from a single founder cell up to more than 2000 cells (Movie 1).

To understand the mechanical processes determining the shape and architecture of biofilms in flow, we developed a 3D multi-scale theoretical framework consisting of two separate models: a cell-based model (Movie 2) and a continuum model (Movie 3). Cells are represented as growing, dividing ellipsoids with pairwise interactions as defined in Ref. [17]; parameters of this model were determined from single-cell biofilm experiments (Table II in SM [24]). Movement of cells occurs through growth and cellular interactions mediated by extracellular matrix and adhesion proteins, but no active motility exists inside *V. cholerae* biofilms. The cell-based model from Ref. [17] was then extended to include flow, as well as previously neglected physical effects that determine biofilm architecture at the single-cell level in strong-flow environments (SM [24]). In particular, each cell feels a force and torque dependent upon its orientation relative to the shear flow; the streamlines of the flow are deformed by the biofilm in a manner consistent with an approximately hemispherical object. Furthermore, the experimental observation that parent and daughter cells adhere to each other at the cell pole for approximately one division time (Fig. 1b) was implemented using Hookean springs, which connect the nearest polar endpoints of these cells and persist for 90% of a division time.

In the complementary continuum model, movement and alignment of biofilm matter is represented through a local mean velocity field ***v*** (*t, **x***) and nematic ‘Q-tensor’ ***Q*** = *S*(***nn*** − 1/3) [12, 13, 28], where *S* (*t, **x***) is the nematic order parameter and ***n***(*t, **x***) is the nematic director field of cellular orientations. Within the biofilm, a modified incompressibility condition, **∇** ⋅ ***v*** = *g*, enforces constant growth with rate *g*; the assumption of a uniform growth rate *g* is valid as long as all cells have access to sufficient nutrients, which holds for the experimental conditions considered here (SM [24]). The effect of this growth, which is directed nematically because cells elongate and divide along their longest axis [29], is imposed by including an additional active term in the stress [30, 31]. Over growth timescales, the passive part of the constitutive relationship can be approximated as purely viscous with effective viscosity *µ*, yielding a stress tensor ***σ*** = −*p***1** + *µ*(**∇*****v*** + **∇*****v**^T^*) − 2*µg**Q*** (SM [24]), where *p* is the pressure. This constitutive law was simulated in the open-source solver Dedalus [32], using a phase-field variable to track the expansion of the biofilm [33, 34]. Thenematic order field was imposed such that ***n*** (*t, **x***) rotates from vertical to an angle just beyond horizontal at the biofilm’s back and sides over a specified length. These rotation lengths in each direction were the key parameters in the model, and were chosen to give quantitative agreement with measured cell alignment fields (Fig. S6). The initial biofilm shape was always taken to be spherically symmetric, so that any asymmetry in the shape was induced by growth along the imposed cell alignment field. Using this model, we could assess to which extent the observed cell alignment fields determined biofilm growth and shape. The combined multi-scale models and experiments revealed that the full growth and cellular alignment program of bacterial biofilms in flow can be categorized into three distinct physical phases (Fig. 1a).

During the initial biofilm growth phase (Fig. 1b,c), the majority of cells are exposed to the flow. We found that the presence of strong shear breaks the otherwise hemispherically-symmetric colony growth, and two key physical processes dominate the cell alignment dynamics. Here we describe and illustrate these two key processes using a combination of scaling laws and the cell-based model. Firstly, daughter cells are reoriented after division to align with the flow by a drag-induced torque caused by the combination of the flow and the polar adhesion to their parent cell (Fig. 1b). Specifically, a horizontal ellipsoidal cell of length *l* and width *r* constrained at one pole, with its longest axis perpendicular to a flow of speed *U*, is expected to feel a torque *τ*_drag_ ∼ *Dl*, where *D* is the drag *D* ∼ *G*_1_*µlU*; here *G*_1_ is a geometric factor [35]. Thus, using 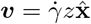, we have 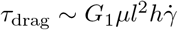, where *h* is the height of the cell centroid from the surface. Secondly, the shear in the *z*-direction applies a torque, causing a cell’s longest axis to rotate about the axis perpendicular to the plane of the flow [36]. For a horizontal cell whose longest axis is parallel to the flow, this torque is approximately 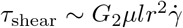; here *G*_2_ is a second geometric factor [37]. Both torques *τ*_drag_ and *τ*_shear_ are expected to be of the same order of magnitude *τ* ∼ 1 pN *µ*m, which is not strong enough to rip fully surface-attached cells from the surface [23]. However, the flow-induced torques act together to cause the verticalization of daughter cells at the front of the biofilm that are not fully surface-attached [38, 39], or that have been partially verticalized by a peeling instability induced by nearby cells [23] (Fig. 1c). Both flow-induced cell reorientation processes were captured by the cell-based model (Fig. 1b,c).

In the second growth phase (Fig. 2), cells in the outer shell of the biofilm are still exposed to the flow, whereas the core of the biofilm is sheltered by surrounding cells and extracellular matrix. We found that the location within the biofilm determines which cell alignment dynamics dominates: cells that are exposed to the flow at the upstream end of the biofilm continue to be realigned vertically owing to the torques *τ*_drag_ and *τ*_shear_ (Fig. 2a,b), whereas cells elsewhere in the outer shell of the biofilm continue to align with the flow, mainly owing to *τ*_drag_, maintaining asymmetric growth of the biofilm overall. In particular, growth of the horizontally-aligned cells in the downstream region causes distinctive droplet-like shapes, which is captured by continuum simulations of growing biofilms with cell alignment fields consisting of a downstream region of flow-aligned cells (Fig. 2a-c). However, cells in the core of the biofilm are not exposed to the flow, so their dynamics are dominated by growth; these cells are subject to a previously observed growth-induced buckling instability [23, 40–42] and the ‘inverse domino’ effect of being surrounded by already vertical cells [23]. We thus conclude that the combination of the flow-induced and growth-induced realignment processes leads to a gradient in the vertical alignment of cells from the upstream end to the downstream end of the biofilm and a wave of cellular verticalization that travels through the biofilm from upstream to downstream (Fig. 2e).

**FIG. 2.**
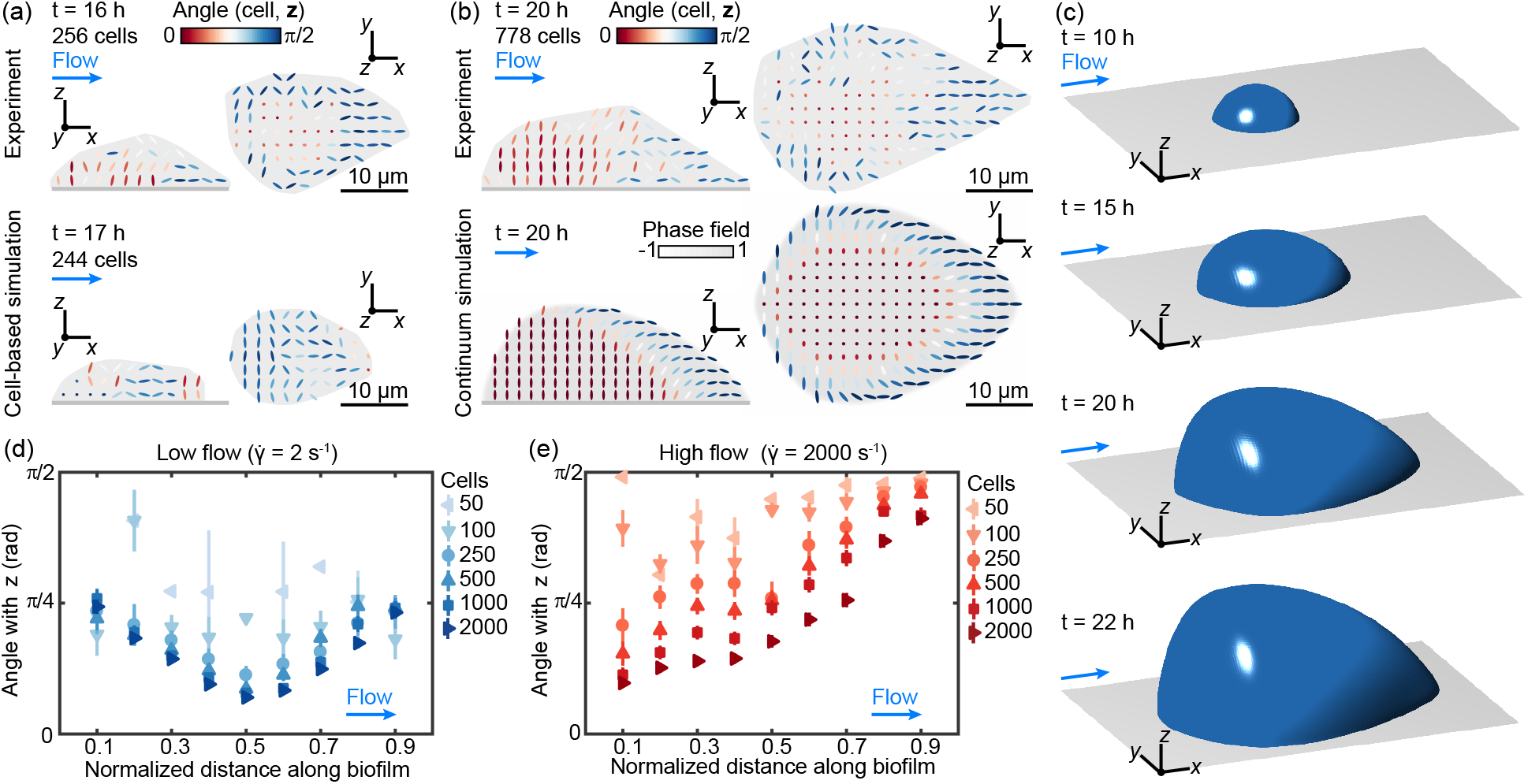
A wave of cell verticalization travels downstream through the biofilm in the second phase of biofilm growth. (a) At the beginning of this phase, a small group of cells is verticalized at the front of the biofilm by a combination of cell-cell interactions and fluid shear. (b) The fraction of vertical cells increases as the biofilm grows. The downstream region of flow-aligned cells leads to distinctive droplet-like shapes that are captured in (a) cell-based (Movie 2) and (b) continuum simulations (Movie 3). For the experiments (*n* = 3) and cell-based simulations (*n* = 10), the grey area denotes the region inside the convex hull around gridpoints with a cell number density higher than 0.1 *µ*m^*−*3^ per biofilm. (c) 3D renderings of shapes generated by the continuum simulations. The isosurface *φ* = 0 of the phase-field variable *φ* is shown (SM [24]). (d) In low flow environments, a central core of the biofilm is verticalized owing to a buckling instability induced by growth and surface attachment. (e) Strong flow causes symmetry breaking and a growing group of vertically aligned cells at the front of the biofilm. Error bars show the standard error for gridpoints spaced 2 *µ*m throughout *n* = 3 biofilms in each case.

In the third growth phase (Fig. 3), the majority of cells in the biofilm are sheltered from the flow, and growth dominates the cell alignment dynamics. Owing to the growth-induced buckling instability and verticalization wave, we observed that biofilms contain a core of highly vertically-aligned cells (Fig. 3a). Sheltered cells in the center of the biofilm have a similar dynamics to those in biofilms in low flow environments, where the shear is not strong enough to reorient cells [17, 19, 23].

**FIG. 3.**
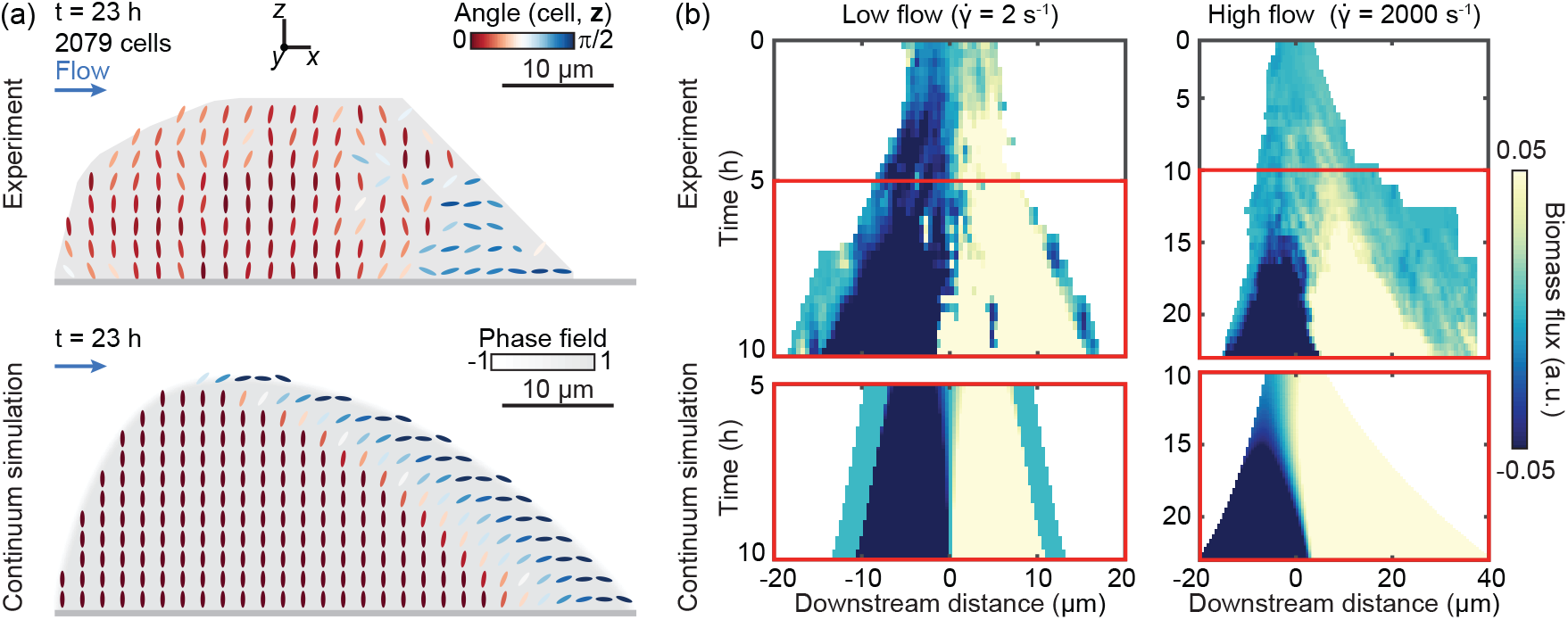
Biofilms transition from asymmetric to symmetric growth in the final phase of their development in flow. (a) In the final phase of growth, most of the cells in the biofilm are vertical, with only a small region of flow-aligned cells on the downstream side. The grey area denotes the region inside the convex hull around gridpoints with a cell number density higher than 0.1 *µ*m^*−*3^ per biofilm (*n* = 3). (b) Growth-induced cumulative biomass flux through the *yz*-plane at each downstream distance; positive and negative values correspond to flux in the downstream and upstream direction, respectively. In low flow, growth is always symmetric (left). In high flow (right), growth symmetry is broken in the early phases, but as more cells verticalize, the biofilm transitions to symmetric growth, which is also captured by continuum simulations (bottom). In experiments (top panels, *n* = 3), biomass flux measurements were obtained using optical flow [17]. In continuum simulations (bottom), the solution for the flow field inside the biofilm was used to calculate cumulative biomass flux.

We used the continuum model, which accounts explicitly for directed cell growth, to investigate how the observed cell alignment fields across the phases determine biofilm growth and shape. We found that in the earlier stages of development, when cells tend to be aligned in a gradient from vertical in the upstream region to horizontal in the downstream region, growth is predominantly in the downstream direction (Fig. 3b, Fig. S8). As the region of vertical cells expands downstream, growth becomes more symmetric, eventually resembling the symmetric radial expansion of biofilms in weak flow (Fig. 3b). In the final phase, growth is preferentially upward owing to the predominance of vertical cells (Fig. S8). The agreement between the growth dynamics observed in our experiments and continuum simulations (Fig. S9) suggests that the competition between flow-aligned and vertical growth is sufficient to explain biofilm growth in flow for biofilms with up to several thousand cells.

In the past, deformation and shear-induced erosion have been hypothesized to explain the flow-induced symmetry breaking of bacterial biofilms in flow [16, 19]. Although this is expected to be true for the extremely large shear rates experienced in turbulent flow, or for bacterial species with weak matrix, we discovered that the asymmetric growth of cells reoriented by the flow is sufficient to account for the architectures of biofilms in our experiments 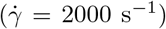. In this physiologically relevant flow regime, shear-induced erosion has been shown to be resisted by the increased production of cell-cell adhesion proteins [17]. Although some cells are still carried away by the flow (Fig. 1c), the effect of erosion is negligible for the architecture dynamics. We now show that deformation is also negligible for biofilms in our experiments. The fluid, which has the viscosity of water *µ*_*w*_, exerts a stress on the biofilm of approximate magnitude 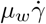 so that, by matching stresses at the fluid-biofilm interface, the strain needed to balance the external stress is approximately 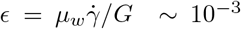, since biofilms have hydrogel-like material properties with elastic modulus *G* ~ 10^3^ Pa [43]. Therefore a *V. cholerae* biofilm will not be significantly deformed by the flow, and over growth time scales, a balance between the internal elasticity and external flow appears instantaneous, with growth then occurring along nematic directions. This supports the hypothesis that nematically-aligned growth is the key determinant of bacterial biofilm shape.

## CONCLUSION

The above experimental and numerical results show that flow initially breaks the symmetry in the cell alignment field of growing biofilms. Because cells grow in the direction of their longest axis, the altered cell orientations significantly affect biofilm architecture, causing distinctive droplet-like shapes. In later stages, cells verticalize in a wave that travels from the upstream end to the downstream end of the biofilm, causing a transition from asymmetric flow-aligned growth to symmetric growth, even in the presence of strong flow. In contrast with previous assumptions, deformation and shear-induced erosion are not important determinants of biofilm architecture for the shear rates studied here. Individual cell dynamics are a crucial for understanding the architecture of growing biofilms, and must be tracked carefully when characterizing the effect of external fields on biofilm growth.

## Acknowledgments

We thank Takuya Ohmura and Vili Heinonen for helpful discussions. Continuum simulations were performed on the *Leavitt* system at the Bates College High Performance Computing Center. This work was partially supported by an Edmund F. Kelly Research Award (J.D.) and James S. McDonnell Foundation Complex Systems Scholar Award (J.D.), as well as the Human Frontier Science Program (K.D., CDA00084/2015-C), the Max Planck Society (K.D.), the European Research Council (K.D., StG-716734), and the Deutsche Forschungsgemeinschaft (SFB 987).

