## Supplemental Material for "Flow-induced symmetry breaking in growing bacterial biofilms"

### Supplemental Material:
### Flow-induced symmetry breaking in growing bacterial biofilms

Philip Pearce,[1] Boya Song,[1] Dominic J. Skinner,[1] Rachel Mok,[1,2] Raimo Hartmann,[3] Praveen K. Singh,[3] Hannah Jeckel,[3] Jeffrey S. Oishi,[1,4] Knut Drescher,[3,5] and Jörn Dunkel[1]

[1]*Department of Mathematics, Massachusetts Institute of Technology, Cambridge MA 02139-4307, USA*
[2]*Department of Mechanical Engineering, Massachusetts Institute of Technology, Cambridge, MA 02139-4307, USA*
[3]*Max Planck Institute for Terrestrial Microbiology, Marburg, Germany*
[4]*Department of Physics, Bates College, Lewiston, ME 04240, USA*
[5]*Department of Physics, Philipps-Universität Marburg, Marburg, Germany*
(Dated: August 15, 2019)

#### SINGLE CELL AGENT-BASED SIMULATIONS

##### Model description

Our single cell model is based on the agent-based framework described in [1], with modifications to include the effect of the flow on the cells and cell-cell polar adhesion. Cells are modeled as ellipsoids of half-length $l$ and half-width $r$; each cell is described by its position $\boldsymbol{x}$, orientation $\hat{\boldsymbol{n}}$ and effective local viscosity $\mu$ . The dynamics of the cells are approximated as over-damped, as cells live at low Reynolds number $Re \approx 10^{-4}$ [1]. Denoting the identity matrix by $\boldsymbol{I}$ and the dynamic viscosity of water by $\mu_w$, the over-damped translational and orientational dynamics for a single cell are

$$\begin{aligned} \frac{\mathrm{d}\boldsymbol{x}}{\mathrm{d}t} &= \boldsymbol{\Gamma}^{-1}\left(-\frac{\partial U_{bdy}}{\partial \boldsymbol{x}} - \frac{\partial V}{\partial \boldsymbol{x}} - \frac{\partial U_s}{\partial \boldsymbol{x}}\right) + \frac{\mu_w}{\mu}\boldsymbol{v}_{flow} \\ \frac{\mathrm{d}\hat{\boldsymbol{n}}}{\mathrm{d}t} &= \left(\boldsymbol{I} - \hat{\boldsymbol{n}}\hat{\boldsymbol{n}}^T\right)\left[\boldsymbol{\Omega}^{-1}\left(-\frac{\partial U_{bdy}}{\partial \hat{\boldsymbol{n}}} - \frac{\partial V}{\partial \hat{\boldsymbol{n}}} - \frac{\partial U_s}{\partial \hat{\boldsymbol{n}}}\right)\right] + \frac{\mu_w}{\mu}\boldsymbol{\omega}_{flow} \end{aligned} \tag{1}$$

where $\boldsymbol{\Gamma}$, $\boldsymbol{\Omega}$ are

$$\begin{aligned} \boldsymbol{\Gamma} &= \gamma_m\left[\gamma_{||}\left(\hat{\boldsymbol{n}}\hat{\boldsymbol{n}}^T\right) + \gamma_{\perp}\left(\boldsymbol{I} - \hat{\boldsymbol{n}}\hat{\boldsymbol{n}}^T\right)\right] \\ \boldsymbol{\Omega} &= \omega_m\omega_R\boldsymbol{I}. \end{aligned}$$

Here, $\gamma_m$ and $\omega_m$ are the typical translational and rotational drag coefficients for Stokes drag in the extracellular matrix for a spheroid ($\gamma_m = 6\pi\mu r$, $\omega_m = 8\pi\mu l r^2$) [1]. $\gamma_{||}$, $\gamma_{\perp}$ and $\omega_R$ are dimensionless geometric parameters characterizing the longitudinal and transverse friction parameters that depend only on the aspect ratio $a = l/r$ of the cell [1].

The pairwise cell-cell interactions are described by two potentials $V$ and $U_s$, where $V$ is the general pairwise cell-cell interaction that applies to all pairs of cells [1], and $U_s$ is the polar attachment that only exists between two sibling cells after division. For two cells $\alpha$ and $\beta$, let $l_\alpha$ and $l_\beta$ be their half-lengths, $r_\alpha$ and $r_\beta$ be their widths, $\hat{\boldsymbol{n}}_\alpha$ and $\hat{\boldsymbol{n}}_\beta$ be their orientation vectors, $r_{\alpha\beta}$ be the distance between their centroids, and $\hat{\boldsymbol{r}}_{\alpha\beta}$ be the unit vector pointing from the centroid of cell $\alpha$ to the centroid of cell $\beta$. The general pairwise cell-cell interaction potential between these two cells, which accounts for short-range cell-cell repulsion due to steric forces, cell-cell repulsion due to osmotic pressure, and cell-cell attraction mediated by adhesion molecules such as RbmA, follows the cell-cell interaction potential formula in [1]

$$U_{\alpha\beta} = \epsilon_0\epsilon_1\left[\nu_{steric}\exp\left(-\frac{\rho_{\alpha\beta}^2}{\lambda_{r,\,steric}^2}\right) + \exp\left(-\frac{\rho_{\alpha\beta}^2}{\lambda_r^2}\right) + \frac{\nu}{1+\exp\left(\frac{\rho_a - \rho_{\alpha\beta}}{\lambda_a}\right)}\right].$$

Here $\rho = \frac{r_{\alpha\beta}}{\sigma}$ is the cell-cell distance normalized by the overlap shape factor $\sigma(l_\alpha, l_\beta, r_\alpha, r_\beta, \hat{\boldsymbol{n}}_\alpha, \hat{\boldsymbol{n}}_\beta, \hat{\boldsymbol{r}}_{\alpha\beta})$ in [1]. The strength of $U_{\alpha\beta}$ is described by $\epsilon_0$ and adjusted by the shape factor $\epsilon_1(l_\alpha, l_\beta, r_\alpha, r_\beta, \hat{\boldsymbol{n}}_\alpha, \hat{\boldsymbol{n}}_\beta)$, depending on the relative cell orientations and cell shapes. $\nu_{steric}$ describes the relative strength of the steric cell-cell repulsion, $\lambda_{r,\,steric}$ describes the range of the steric cell-cell repulsion, $\lambda_r$ describes the range of the osmotic cell-cell repulsion, $\nu$ describes the relative strength of cell-cell attraction, $\rho_a$ describes the attraction position, and $\lambda_a$ describes the width of the cell-cell attraction. The cell-cell potential also has an associated translational relaxation time $\tau_t$, which can be interpreted as a time scale of how long it takes for a bacterium to reach an equilibrium configuration from the cell-cell interaction potential. We used the same values as in [1] for the model parameters $\lambda_{r,\,steric}$, $\lambda_r$, $\nu$, $\rho_a$ and $\lambda_a$. The value of $\epsilon_0$ was increased compared with [1], in order to account for the increase of the cell-flow interaction energy $\epsilon_{flow}$ in the simulations with increased flow; therefore $\tau_t$ was slightly decreased. We found that choices of $\epsilon_0 = 10^4\epsilon_{flow}$, $\nu_{steric} = 1$ and $\tau_t = 5.65$ s were sufficient to ensure the cells do not overlap in these simulations; the osmotic and the short-range steric parts of the repulsion were both necessary for this. All the interaction potential parameter values are listed in Table II. For a single cell $\alpha$ in a biofilm with $N$ cells, $V$ is the total potential for all $N-1$ pairwise cell-cell interactions between cell $\alpha$ and other $N-1$ cells $\beta$ ($V = \sum_{\beta=1,\beta\neq\alpha}^{N} U_{\alpha\beta}$).

We model the polar cell-cell attachment after division using a harmonic spring between two sibling cells with spring constant $k_s$ and natural length $r_s$. As soon as a cell divides, a spring is assigned between the two closest endpoints of the daughter cells, with the spring potential

$$U_s = k_s \left(r_{endpoint} - r_s\right)^2 ,$$

where $r_{endpoint}$ is the distance between the two closest endpoints. To match with experiments, where polar cell-cell adhesion was observed to last for approximately one division time, the spring breaks at $0.9 \cdot \tau_g$ after division, where $\tau_g$ is the average doubling time measured from single-cell experimental data using the same technique as [1]. The natural spring length $r_s$ was chosen to match the cell half-width for simplicity. The spring constant $k_s$ was chosen to be large enough such that cells are not carried away by the flow immediately after division in the initial stages of biofilm growth. The values of $\epsilon_s$, $r_s$, $k_s$ and $\tau_g$ are listed in Table II.

The interaction between cells and the wall boundary is modeled with the same repulsive interaction potential as in [1]

$$U_{bdy} = \begin{cases} 0 & z_o \leq 0 \\ \epsilon_{bdy} \exp\left(\frac{z_o}{\sigma_{bdy}}\right) & z_o > 0 \end{cases}$$

where $\epsilon_{bdy}$ captures the magnitude of the cell-boundary interaction, and $\sigma_{bdy}$ captures the range of the interaction. $z_o$ is an overlap coordinate defined as $z_o = l|\hat{\boldsymbol{n}} \cdot \hat{\boldsymbol{N}}| + r - \hat{\boldsymbol{N}} \cdot (\boldsymbol{x} - \boldsymbol{S})$ where $\hat{N}$ is the normal vector of the plane, and $\boldsymbol{S}$ is any point on the plane. Here we use $\hat{N} = [0, 0, 1]$ and $\boldsymbol{S} = [0, 0, 0]$ such that the wall is the xy-plane that crosses the origin. The values of $\epsilon_r = \epsilon_{bdy}/\epsilon_0$ and $\sigma_{bdy}$, listed in Table II, were chosen to be the same as the values in [1].

The instantaneous cell length growth follows the growth equation in [1]

$$\frac{\mathrm{d}l}{\mathrm{d}t} = \frac{l}{\tau_g} \ln(2) \tag{2}$$

where $l$ is the instantaneous half-length of the cell and $\tau_g$ is the average doubling time. Cell widths are constant throughout simulations. A cell divides when it grows an additional length $\Delta l$ from its birth length, where $\Delta l$ is drawn from a Gaussian distribution with mean $l_{add}$ and standard division $\sigma_{add}$. Cells divide into two daughter cells that have half the length of their parent cells and the same orientation as their parent cell. The values of $l_{add}$ and $l_{add}$ were fitted according to the experimental distribution of cell lengths. Both values are listed in Table II.

To account for the increase in the local effective viscosity of a cell as extracellular polymeric substances (EPS) matrix and cell-cell adhesion proteins are produced by itself and the surrounding cells, we used two steps to calculate the local viscosity $\mu$ experienced by each cell. First, to account for extracellular matrix and adhesion protein generation by each cell, the individual local viscosity contribution of a cell is a sigmoid function that increases from $\mu_w$ to $\mu_{max}$, where $\mu_w$ is the viscosity of water. Let $t_{age}$ be the cell's age, scaled by the cell's division time. The individual local viscosity contribution of the cell is

$$\mu_{indv} = \mu_{max} - \frac{\mu_{max} - \mu_w}{1 + \exp\left(\frac{t_{age} - t_0}{\Delta t}\right)}$$

where $t_0 = 0.4$ is the viscosity transitional time and $\Delta t = 0.1$ is the viscosity-increase time scale (both are a proportion of the cell's age). The left panel of Figure S1 shows a plot of of $\mu_{indv}$ as a function of $t_{age}$. Then, to account for extracellular matrix and adhesion proteins generated by neighboring cells, the local viscosity experienced by a cell $\alpha$ is taken to be the Gaussian-filtered value of the individual local viscosity contributions of the surrounding cells

$$\mu_\alpha = \sum_{\beta=1}^{N} \frac{1}{\sqrt{\pi}\sigma_{vis}} \exp\left(-\frac{r_{\alpha\beta}^2}{\sigma_{vis}^2}\right) \mu_{indv,\,\beta},$$

where $\sigma_{vis}$ is the viscosity averaged length scale and $r_{\alpha\beta}$ is the distance between the centroids of cell $\alpha$ and cell $\beta$. The right panel of Figure S1 shows the local viscosity distribution of a simulated biofilm with 250 cells. The viscosity parameters $\mu_{max} = 2 \cdot 10^5$ Pas and $\sigma_{vis} = 1.39$ µm were chosen by matching the translational and rotational dynamics of cells aligning with the flow in simulated biofilms with those of experimental ones (Fig. S2). The value of $\mu_{max}$ is listed in Table II.

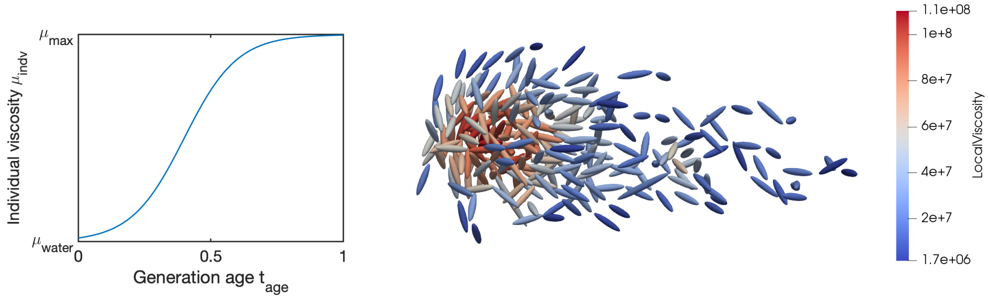


FIG. S1. Left: $\mu_{indv}$ as a function of cell generation age $t_{age}$. Right: The top view of a simulated biofilm with 250 cells, where cells are colored by their local viscosity.

When the biofilm is equal to or smaller than 20 cells, we approximate the flow field as the linear shear flow $\boldsymbol{v}_{flow} = \dot{\gamma} z \hat{\boldsymbol{x}}$, where $z$ is the height of the cell above the surface and $\dot{\gamma}$ is the constant shear rate of the linear shear flow. When the biofilm is larger than 20 cells, we approximate the flow field around the biofilm by a linear shear flow passing over a hemispherical bump on a plane floor. The center of the hemispherical bump is located at the $xy$-center of the biofilm, and the radius of the hemispherical bump is approximated as the 25th percentile of distances between all the cells to the $xy$-center of the biofilm. Let $a$ be the radius of the hemispherical bump, and $(x_c, y_c, z_c)$ be the position of the center of the hemispherical bump in Cartesian coordinates. For a point $(x, y, z)$ in Cartesian coordinates, define spherical polar coordinates $(r, \theta, \phi)$ by

$$\frac{x - x_c}{a} = r \sin\theta \cos\phi, \quad \frac{y - y_c}{a} = r \sin\theta \sin\phi, \quad \frac{z - z_c}{a} = r \cos\theta$$

so that the boundary of the hemisphere is given by $r = 1, \theta \in [0, \pi/2]$, and the planar floor is given by

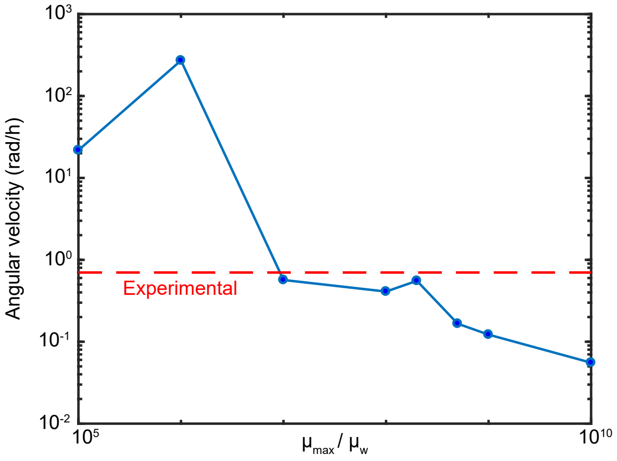


FIG. S2. Average angular velocity of simulated cells aligning to the flow direction from their orientation at birth as a function of maximum viscosity $\mu_{max}/\mu_w$, where $\mu_w$ is the viscosity of water. The value $\mu_{max} = 2 \cdot 10^8 \mu_w$ was chosen to recreate the average angular velocity of flow-aligning cells measured in experiments (0.7 rad/h, dashed red line), while ensuring cells are not carried away by the flow.

$r \geq 1, \theta = \pi/2$. From [2], the flow velocity $\boldsymbol{v}_{flow}$ at $(x, y, z)$ is

$$\begin{aligned}
\frac{\boldsymbol{v}_{flow}}{\dot{\gamma}} =& z\hat{\boldsymbol{x}} - \left( u\cos\phi\hat{\boldsymbol{r}} + v\cos\phi\hat{\boldsymbol{\theta}} + w\sin\phi\hat{\boldsymbol{\phi}} \right) \\
=& \left[ z - (u\sin\theta + v\cos\theta)\cos^2\phi + w\sin^2\phi \right] \hat{\boldsymbol{x}} \\
& - (u\sin\theta + v\cos\theta + w)\cos\phi\sin\phi\hat{\boldsymbol{y}} \\
& + (-u\cos\theta\cos\phi + v\sin\theta\cos\phi)\,\hat{\boldsymbol{z}}
\end{aligned}$$

where $u$, $v$, $w$ are functions of $r$ and $\theta$:

$$\begin{aligned}
u(r,\theta) &= \cos\theta W(r,\cos\theta) + \frac{1}{2}\sin\theta\left[U(r,\cos\theta) + V(r,\cos\theta)\right] \\
v(r,\theta) &= -\sin\theta W(r,\cos\theta) + \frac{1}{2}\cos\theta\left[U(r,\cos\theta) + V(r,\cos\theta)\right] \\
w(r,\theta) &= \frac{U(r,\cos\theta) - V(r,\cos\theta)}{2}.
\end{aligned}$$

$U$, $V$ and $W$ are functions of $r$ and $\cos\theta$. Let $P_m^n$ denote an associated Legendre function in $\cos\theta$ of degree $n$ and order $m$. The expressions of $U$, $V$ and $W$ from [2] are:

$$\begin{aligned}
W(r,\cos\theta) &= \frac{1}{2}\sum_{n=1}^{\infty}\frac{1}{r^{2n+1}}\left[A_{2n+1}\left(\cos\theta P_{2n+1}^1 + \frac{P_{2n}^1}{2n}\right) + rA_{2n}\cos\theta P_{2n}^1\right] \\
U(r,\cos\theta) &= \frac{1}{2}\sum_{n=1}^{\infty}\frac{P_{2n+1}^2}{r^{2n+1}}\left(\frac{A_{2n+1}\cos\theta}{2n} + \frac{rA_{2n}}{4n+1}\right) \\
&\quad + \frac{1}{2}\sum_{n=2}^{\infty}\frac{P_{2n-1}^2}{(n-1)(2n-1)r^{2n}}\left[\frac{(2n-3)(2n+1)A_{2n}}{4n+1} + G_{2n}\right] \\
V(r,\cos\theta) &= -\frac{1}{2}\sum_{n=1}^{\infty}\frac{(2n+1)P_{2n+1}^0}{r^{2n+1}}\left(A_{2n+1}\cos\theta + \frac{2nrA_{2n}}{4n+1}\right) + \sum_{n=1}^{\infty}\frac{G_{2n}P_{2n-1}^0}{r^{2n}}.
\end{aligned} \tag{3}$$

We followed [2] to take the first fifteen terms in Eqs. (3) with the coefficients recorded in Table I.

| $n$ | $A_{2n+1}$ | $A_{2n}$ | $G_{2n}$ |
|---|---|---|---|
| 1 | -2.04530 | 2.75543 | 1.65326 |
| 2 | -0.86783 | 1.82574 | -0.73104 |
| 3 | 0.14321 | 0.08434 | -0.06369 |
| 4 | -0.04430 | -0.04420 | 0.05474 |
| 5 | 0.01643 | 0.02520 | -0.04337 |
| 6 | -0.00640 | -0.01535 | 0.03381 |
| 7 | 0.00232 | 0.00985 | -0.02643 |
| 8 | -0.00054 | -0.00659 | 0.02083 |
| 9 | -0.00024 | 0.00455 | -0.01657 |
| 10 | 0.00057 | -0.00323 | 0.01331 |
| 11 | -0.00069 | 0.00235 | -0.01079 |
| 12 | 0.00070 | -0.00174 | 0.00881 |
| 13 | -0.00067 | 0.00131 | -0.00725 |
| 14 | 0.00062 | -0.00100 | 0.00601 |
| 15 | -0.00056 | 0.00077 | -0.00500 |

TABLE I. First fifteen of $\{A_{2n+1}, A_{2n}, G_{2n}\}$ in Eqs. (3) [2].

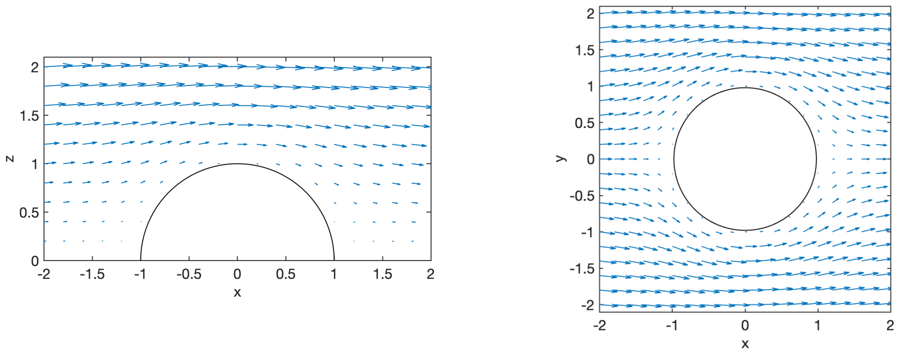


FIG. S3. $\boldsymbol{v}_{flow}/\dot{\gamma}$ for $y = 0$ on the xz-plane (left) and for $z = 0.2$ on the xy-plane (right).

Let $a = l/r$ be the aspect ratio of a bacterium. The rotation caused by the shear flow is computed according to [3]

$$\boldsymbol{\omega}_{flow} = (\boldsymbol{I} - \hat{\boldsymbol{n}}\hat{\boldsymbol{n}}^T)\left[\frac{a^2-1}{a^2+1}\frac{\nabla\boldsymbol{v}_{flow} + \nabla\boldsymbol{v}_{flow}^T}{2} + \frac{\nabla\boldsymbol{v}_{flow} - \nabla\boldsymbol{v}_{flow}^T}{2}\right]\hat{\boldsymbol{n}}.$$

The flow velocity gradient $\nabla\boldsymbol{v}_{flow}$ is computed using a central difference scheme with $\Delta x = \Delta y = \Delta z = 0.01r$.

The key parameters used for the simulations are shown in Table II.

| Parameter | Value | Unit | Description |
|---|---|---|---|
| $r$ | 0.2775 | µm | Average half-width of the bacteria [1]. |
| $\tau_g$ | 8630 | s | Growth time constant (average cell division time of biofilm-associated cells in strong flow, $\dot{\gamma} = 2000\ \mathrm{s}^{-1}$, obtained from experiments). |
| $\dot{\gamma}$ | 2000 | $\mathrm{s}^{-1}$ | Experimental shear rate. |
| $\mu_w$ | 1 | mPa·s | Dynamic viscosity of water at room temperature. |
| $\mu_{max}$ | $2{\cdot}10^5$ | Pa·s | Estimate of the maximal effective dynamic viscosity, owing to extracellular matrix and adhesion proteins, at room temperature. |
| $\epsilon_r$ | 10 | | Ratio comparing the strength of bacteria-boundary interaction to the strength of the bacteria-bacteria interaction ($\epsilon_r = \epsilon_{bdy}/\epsilon_0$) [1]. |
| $\sigma_{bdy}$ | 0.2775 | µm | Non-dimensional boundary potential length scale parameter [1]. |
| $\tau_t$ | 5.65 | s | Translational time scale due to repulsion in matrix (typical time needed for daughter cells in matrix to reach their equilibrium configurations due to repulsion after cell division). |
| $l_{add}$ | 1.01 | µm | Average value of length added to bacteria after division to compute division length. |
| $\sigma_{add}$ | 0.08 | µm | Standard deviation of length added to bacteria after division to compute division length. |
| $\epsilon_0$ | $5{\cdot}10^{-14}$ | J | Strength of the osmotic pressure-mediated cell-cell repulsion. |
| $\nu_{steric}$ | 1 | | Relative strength of the steric cell-cell repulsion. |
| $\lambda_{r,\,steric}$ | 0.83 | | Range of the steric cell-cell repulsion (corresponds to 0.58 µm at a typical overlap factor of $\sigma = 0.7$µm, which is the value it would take for a sphere with the typical mean cell volume of 0.4µm$^3$) [1]. |
| $\lambda_r$ | 1.65 | | Range of the osmotic pressure-mediated cell-cell repulsion (corresponds to 1.16 µm at a typical overlap factor of $\sigma = 0.7$µm) [1]. |
| $\nu$ | 0.13 | | Relative strength of the attractive part of the cell-cell potential [1]. |
| $\lambda_a$ | 0.16 | | Well-width of the attractive part of the cell-cell potential [1]. |
| $\rho_a$ | 2.93 | | Position of the attractive part of the cell-cell potential [1]. |
| $k_s$ | $6.5{\cdot}10^{-1}$ | J·µm$^{-2}$ | Spring constant for the additional directional cell-cell attraction. |
| $r_s$ | 0.2775 | µm | Natural length of the spring between sibling cells. |

TABLE II. Simulation parameters.

**Model implementation**

A custom, highly parallelized individual cell-based code employing graphics processing units (GPUs) was developed to perform the simulations based on [1]. At each time step, cell-cell interactions between all pairs of cells are evaluated. A standard explicit Euler scheme is used to numerically integrate Eqs. (1) and Eq. (2) in non-dimensional form, with $r = 0.2775$µm as the length scale [1], the translational time $\tau_t = 5.65$s as the time scale [1] and $\epsilon = 5 \cdot 10^{-20}$J [1] as the energy scale.

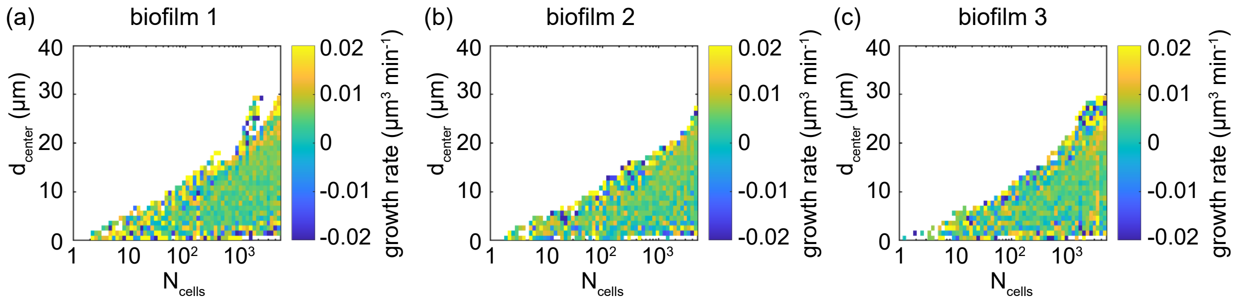


FIG. S4. Heatmaps showing spatially resolved single-cell measurements of growth rate inside three different biofilms (a)-(c) at $\dot{\gamma} = 2000\mathrm{s}^{-1}$ [1]. The growth rate is spatially uniform throughout development.

#### CONTINUUM SIMULATIONS

##### Model description

A biofilm is a mixture of several phases, including cells, extracellular polymeric substances (EPS) and adhesion proteins. In addition to mechanical phases, there are also other relevant fields such as a nematic order field describing cell orientations and nutrient concentration fields. Multi-phase continuum equations for growing biofilms have been postulated [4], which can describe inhomogeneous biofilms with spatial nutrient gradients. However, during the early stages of biofilm growth, the nutrient concentration is approximately constant, as is the growth rate (Fig. S4) [1]. We assume that when the continuum model is applicable, the relative volume fraction of cells and extracellular matrix stays approximately constant, and that the nutrients needed for growth diffuse into the biofilm in a way that does not effect the mechanics. These assumptions are only valid during the early stages of growth, but allow us to treat the biofilm mechanically as a single phase system. The specific form of the mechanics is then determined by the constitutive relation for the stress tensor $\boldsymbol{\sigma}$. We take the system as growing, but otherwise incompressible, i.e. the biofilm grows but the density does not significantly fluctuate, an assumption that is valid only in the early stages of biofilm development. In this case, we must solve a modified incompressibility condition

$$\nabla \cdot \boldsymbol{v} = g.$$

Neglecting inertia, the momentum balance takes the form,

$$\nabla \cdot \boldsymbol{\sigma}^{dev} = \nabla p,$$

where $\boldsymbol{\sigma}^{dev}$ is the deviatoric part of the stress tensor. All that remains is to choose a constitutive law for $\boldsymbol{\sigma}^{dev}$ which accounts for both passive and growth stresses. A full constitutive model would account for the viscoelastic matrix component as well as the nemato-elasticity conferred by the nematic ordering of the cells. For the minimal continuum model we seek a simplified constitutive law that describes the long time growth of the biofilm along nematic directions

The extracellular matrix is known to behave viscoelastically, [4] however, a viscoelastic liquid behaves as a viscous fluid over time scales longer than the relaxation time scale. Since growth occurs over relatively long time scales, we assume that the extracellular matrix simply contributes a viscous part to the constitutive equation.

Biofilms are living liquid crystals and have an internal nematic order [1, 5]. We introduce a nematic order parameter $\boldsymbol{Q}$ measuring the local alignment of the cells [6, 7]. Aside from growth, there are two contributions to the constitutive relation. First, there is a free energy associated with the alignment of the cells; thus any distortion away from an aligned state has an energetic cost, and causes a stress to be exerted on the material. Second, there is anisotropic dissipation depending on whether the macroscopic flow is aligned with the nematic director or not, and dissipation when the director field rotates relative to the fluid.

To create a minimal continuum model, we neglect any anisotropy in the passive part of the stress tensor, reducing the constitutive equation to that of a Newtonian fluid, $\boldsymbol{\sigma}_p = 2\mu \boldsymbol{D}$, where $\boldsymbol{D} = \frac{1}{2}(\nabla \boldsymbol{v} + \nabla \boldsymbol{v}^T)$. We neglect the external fluid shear stress experienced by the biofilm in order to examine the mechanism of growth along nematic directions together with local flow-induced realignment of cells. Order of magnitude estimates (see main text) show that the external shear is insufficient to account for any noticible change in the shape, so we take the external fluid as initially at rest, and of a low relative viscosity ($\mu_{ext} = 0.1\mu$). We also assume that $g$ is not a function of the pressure or the local alignment strength; each cell grows and divides at a constant rate independent of the nematic ordering, in agreement with experimental measurements (Fig. S4).

To account for growth in our consitutive law, we consider the additional stress caused by directed growth. A growing cell with orientation $\boldsymbol{n}$, and position $\boldsymbol{y}$, exerts a force dipole on the surrounding medium,

$$\boldsymbol{f}(\boldsymbol{x}; \boldsymbol{y}, \boldsymbol{n}) = f\delta(\boldsymbol{y} + \ell \boldsymbol{n} - \boldsymbol{x})\boldsymbol{n} - f\delta(\boldsymbol{y} - \boldsymbol{x})\boldsymbol{n} \approx -f\ell \boldsymbol{n}\boldsymbol{n} \cdot \nabla_{\boldsymbol{x}} \delta(\boldsymbol{x} - \boldsymbol{y})$$

To find the additional contribution to the stress tensor due to such dipoles, we use Kirkwood's formula [8],

$$\boldsymbol{\sigma}_g = -f\ell\rho\langle \boldsymbol{n}\boldsymbol{n}\rangle = -\zeta g \langle \boldsymbol{n}\boldsymbol{n}\rangle = -\zeta g \left(\boldsymbol{Q} + \frac{1}{d}\boldsymbol{1}\right).$$

where $\boldsymbol{Q} = \langle \boldsymbol{n}\boldsymbol{n}\rangle - \boldsymbol{1}/d$ is the nematic order parameter, $\zeta > 0$ is a constant, $d$ is the dimension (all simulations were performed with $d = 3$), and we have assumed the strength of the dipole is proportional to the growth rate. The isotropic part of this stress can be absorbed into the pressure; note, however, that the pressure is a Lagrange multiplier for $\nabla \cdot \boldsymbol{v} = g$ and hence accounts for growth.

Since $\zeta g = f\ell\rho$ is written in terms of microscopic quantities, its value could in theory be measured. Here, we take a phenomenological approach. Consider a 3D element of biofilm that is surrounded by a zero viscosity fluid at zero pressure. Aligning the nematic axis with $\hat{z}$, we have

$$\boldsymbol{v} = \left(\frac{g-a}{2}x, \frac{g-a}{2}y, az\right),$$

for some $0 < a \leq g$. The combined stress tensor $\boldsymbol{\sigma}^{dev} = \boldsymbol{\sigma}_p^{dev} + \boldsymbol{\sigma}_a^{dev} = -\zeta g \boldsymbol{Q} + 2\mu \boldsymbol{D}$ is diagonal, with

$$\begin{aligned} \sigma_{xx}^{dev} &= \sigma_{yy}^{dev} = \mu(g-a) + \zeta g S/3 \\ \sigma_{zz}^{dev} &= 2\mu a - 2\zeta g S/3, \end{aligned}$$

where $Q_{xx} = Q_{yy} = -S/3$, $Q_{zz} = 2S/3$, and $S$ is the strength of alignment. We suppose that for $S = 1$, i.e. perfect alignment in the $\hat{z}$ direction, the only motion is in the $\hat{z}$ direction. This is possible only for $\zeta = 2\mu$. A similar calculation in 2-dimensions requires $\zeta = \mu$, so

$$\zeta = (d-1)\mu.$$

The full continuum equations inside the growing biofilm can be written as

$$\begin{aligned} \nabla \cdot \boldsymbol{v} &= g \\ \nabla \cdot \boldsymbol{\sigma}^{dev} &= \nabla p \\ \boldsymbol{\sigma}^{dev} &= 2\mu \boldsymbol{D} - (d-1)\mu g \boldsymbol{Q}. \end{aligned}$$

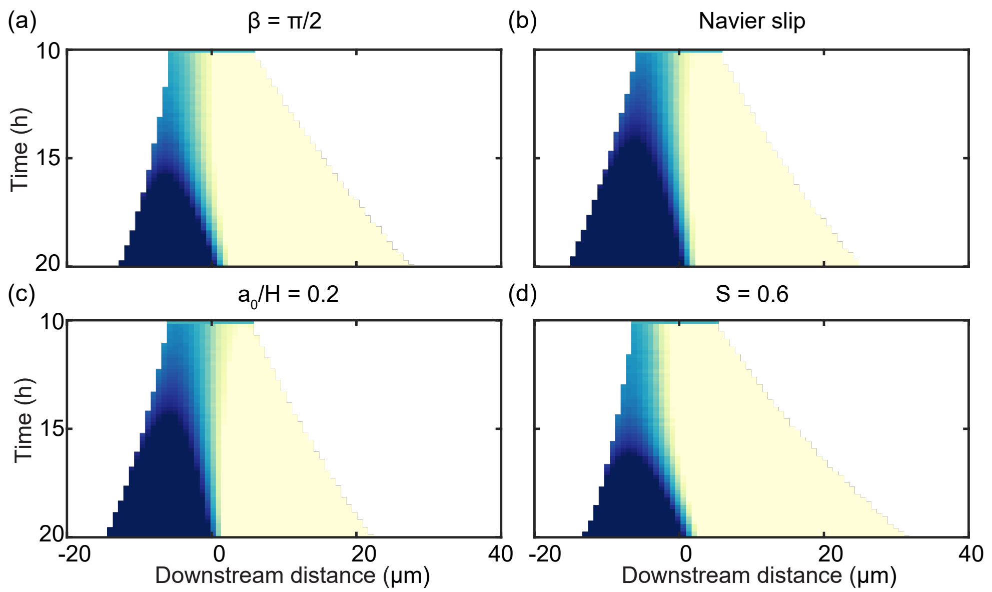


FIG. S5. The continuum model's prediction of a transition from asymmetric to symmetric growth is not sensitive on the model parameters, and is recreated robustly for (a) downstream angle $\beta = \pi/2$, (b) a standard Navier slip condition, (c) director field transition length at the back of the biofilm $a_0/H = 0.2$ and (d) nematic order parameter $S = 0.6$. The values of these parameters used in the main paper are summarized in Table III.

##### Boundary conditions

This minimal continuum model represents growth along an imposed nematic direction. In order to simulate the model, we must impose boundary conditions at the biofilm-fluid, and biofilm-surface interface. We take the external fluid as initially at rest, at which point the phase-field method automatically enforces continuity at the internal interface.

At the biofilm-surface interface we impose a modified Navier-slip condition, which can be written as

$$\mu \frac{\partial u}{\partial z} = e_s(1 - (\boldsymbol{n} \times \hat{z})^2)u,$$
$$\mu \frac{\partial v}{\partial z} = e_s(1 - (\boldsymbol{n} \times \hat{z})^2)v,$$
$$w = 0,$$

where $\boldsymbol{v} = (u, v, w)$, we have assumed the surface is defined by $z = 0$, and $e_s$ is a slip parameter. The modification from a regular Navier-slip is the inclusion of the $(1 - (\boldsymbol{n} \times \hat{z})^2)$, where $\boldsymbol{n}$ is the nematic director. This is to reflect the fact that horizontal cells continue to grow, leading to biofilm movement along the surface in areas with many horizontal cells, whereas vertical cells are fully attached to the surface, resisting movement along the surface. We found that simulations with a regular Navier-slip condition did not produce qualitatively different results (Fig. S5).

##### Imposed nematic field

In our minimal continuum model, we do not directly solve for the nematic field, but impose it based on experimental data (see Fig. 2e in the main text). Physically, the director field is set by interactions between the cells and the flow, the cells and the wall, and cell-cell interactions. Verticalization of cells at the base of

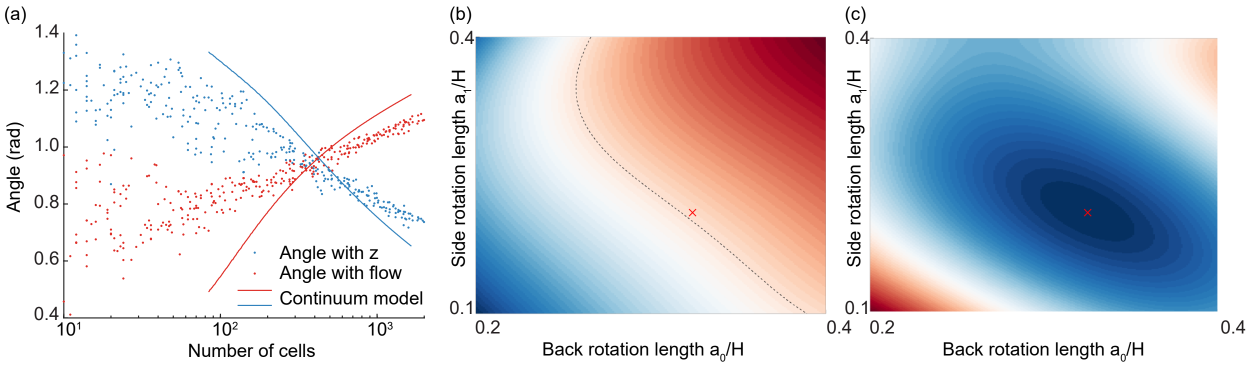


FIG. S6. Key governing parameters in the continuum model and quantitative justification for their values. (a) Average angle with $z$ (blue) or the flow direction (red) versus number of cells for experiments (dots) and continuum simulations with chosen values of the key parameters $a_0$ and $a_1$ (lines). The time variable in continuum simulations was converted to number of cells by fitting an exponential curve to time versus number of cells for $n = 3$ biofilms and rearranging the obtained equation. (b) Dependence of the time of transition between phases 2 and 3, as defined in Fig. 1a, on the parameters $a_0$ and $a_1$ in the continuum model. The line shows the values for which the transition time is 17h, which is approximately the value found in experiments. Note that a transition still occurs for a range of values of $a_0$ and $a_1$, showing that the choice of their values does not qualitatively affect the results (see also Fig. S5). (c) Mean-squared distance (MSD) between average angle curves shown in (a) and experimental values. The values of $a_0$ and $a_1$ were taken to be the values that minimized the MSD. The heatmaps in (b) and (c) were generated by performing 24 simulations and interpolating between them. The red crosses in (b) and (c) show the chosen values of $a_0$ and $a_1$.

the biofilm quickly propagates into the interior, with only a boundary layer of cells at the edge rotated by the flow. We seek a minimal continuum model to investigate how growth along nematic directions affects the shape of the biofilm without being a full mechanical model accounting for all stresses and strains of the system. Therefore we impose the nematic order based on experimental measurements, acknowledging that this is set by interactions at the single cell scale that our minimal continuum model does not account for.

We take the nematic director as $\boldsymbol{n} = (n_x, n_y, n_z) = (\cos\alpha_0, \sin\alpha_0 \sin\alpha_1, \sin\alpha_0\cos\alpha_1)$. The angle $\alpha_0$ is calculated based on the relative distance to the back of the biofilm; given a point $(x, y, z)$ within the biofilm, the back defined as $(x^*, y, z)$, with $x^* = \max_s\{s|(s, y, z) \text{ is in biofilm}\}$. The angle $\alpha_0$ changes linearly over a distance $a_0$ from the back, so that

$$\alpha_0 = \min\left\{\frac{\pi}{2} - \beta\frac{x - x^* + a_0}{a_0},\ \frac{\pi}{2}\right\} \tag{4}$$

where $\beta$ is the angle through which the director field is rotated. A value of $\beta = 1.25 \cdot \frac{\pi}{2}$ was used in simulations, so the director rotates from being vertical at the front of the biofilm, to being slightly beyond horizontal at the back, as observed in experiments (see Fig. 3 in the main paper). The rotation through $\alpha_1$ ensures that the director rotates outwards, away from the central plane of the biofilm, defined by $y = y_c$. The angle $\alpha_1$ is defined similarly to $\alpha_0$, but calculated based on the distance to either the left or the right side of the biofilm. Defining $y_+$ $(y_-) = \max_s\,(\min_s)\{s|(x, s, z) \text{ is in biofilm}\}$, we take

$$\alpha_+ = \max\left\{\frac{\pi}{2}\frac{(y - y_+ + a_1)}{a_1}, 0\right\}, \qquad \alpha_- = \min\left\{-\frac{\pi}{2}\frac{(y - y_- + a_1)}{a_1}, 0\right\},$$

$$\alpha_1 = \alpha_+ + \alpha_-.$$

We found that the numerical results did not depend qualitatively on the parameters $a_0$, $a_1$, or $\beta$; a transition from asymmetric to symmetric growth occured for a wide range of values of these parameters (Fig. S5). However, the choices of $a_0$ and $a_1$ were the key quantitative control parameters in the model that determined when the transition occured (Fig. S6).

#### Numerical phase-field method

To solve for the biofilm growth numerically we use the phase-field method for complex fluids [9]. We introduce an order parameter $\phi$, where $\phi = +1$ in the biofilm and $\phi = -1$ outside. When $\phi = \pm 1$ the equations become the governing equations of each phase. The only regions where this is not true are the interfacial regions. The width of the interfacial region is set by the choice of mixing energy, which penalizes interfaces whilst encouraging demixing. By reducing the size of the interface one can take a limit of the phase-field equations to formally recover the sharp interface equations with correct surface boundary conditions. Thus, by solving the phase field equations, with a small interface region, one can approximately solve the full problem without needing to explicitly keep track of the internal interfaces.

Our equations only differ from [9], in that in our formulation $\nabla \cdot \boldsymbol{v} \neq 0$, and consequently an additional term $g(1-\phi^2)/2$ appears in the Cahn-Hilliard equation, representing growth in interfacial regions. The phase field equations are,

$$
\begin{aligned}
\nabla \cdot \boldsymbol{v} &= \frac{1+\phi}{2} g, \\
0 &= -\nabla p + \nabla \cdot, \boldsymbol{T}, \\
\boldsymbol{T} &= \frac{1+\phi}{2} \boldsymbol{\sigma}_b^{dev} + \frac{1-\phi}{2} \boldsymbol{\sigma}_w - \lambda \nabla \phi \otimes \nabla \phi, \\
\phi_t + \boldsymbol{v} \cdot \nabla \phi &= \frac{g(1-\phi^2)}{2} + \gamma_1 \lambda \nabla^2 \left[ -\nabla^2 \phi + \frac{\phi(\phi^2 - 1)}{\epsilon^2} \right], \\
\boldsymbol{\sigma}_{ext} &= 2\mu_{ext} \boldsymbol{D}, \\
\boldsymbol{\sigma}_b^{dev} &= 2\mu \boldsymbol{D} - (d-1)\mu g \boldsymbol{Q},
\end{aligned}
$$

where $\epsilon$ controls the width of the interface, $\lambda$ controls the strength of surface tension and is set so the effective capillary number is large. A large capillary number ensures that the system is not dominated by surface tension, so that the biofilm can take non-hemispherical shapes; here it is important to reiterate that deformation by the imposed flow is not a key driving process of the biofilm dynamics (as discussed in the main text), and is therefore not accounted for in our continuum model. A large capillary number also means that the contact angle condition does not affect the numerical solution. $\gamma_1$ sets the time scale for phase field diffusion, and $\mu_w$ is the viscosity of water. All parameters used are listed in Table III. Boundary conditions on $\phi$ are given by

$$
\begin{aligned}
\boldsymbol{m} \cdot \nabla \phi &= 0, \\
\boldsymbol{m} \cdot \nabla \nabla^2 \phi &= 0,
\end{aligned}
$$

for a boundary with normal $\boldsymbol{m}$, which, when combined, represent no flux and a contact angle of $90^o$ [9].

We solve the equations in a channel of height $H$, width $1.2H$, and length $4H$. As long as $H$ is large enough to contain the biofilm, the choice of $H$ does not affect the solution. We non-dimensionalize the equations using $H$ as a length scale, $1/g$ as a time scale (so that $\boldsymbol{v} = Hg\boldsymbol{v}^*$, where $\boldsymbol{v}^*$ is dimensionless) and similar for

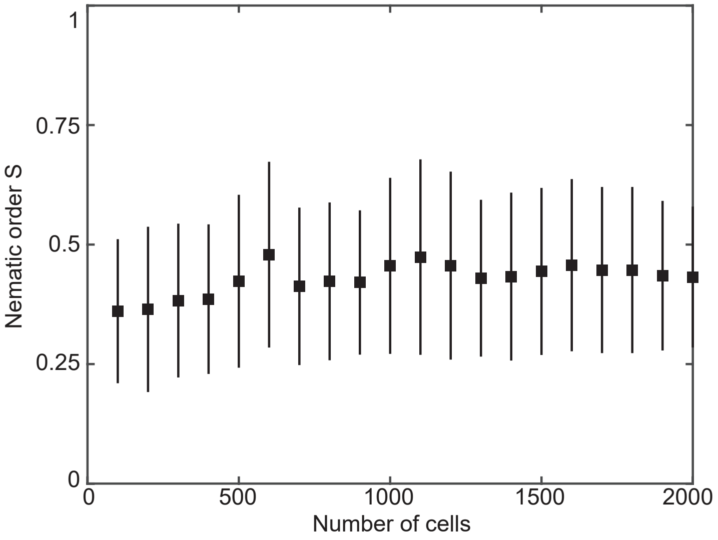


FIG. S7. Mean nematic order parameter $S$ at gridpoints spaced 2 $\mu$m throughout $n = 3$ biofilms. Error bars show the standard deviation.

other variables. The equations become

$$\begin{aligned}
\nabla^* \cdot \boldsymbol{v}^* &= \frac{1+\phi}{2} g^*, \\
0 &= -\nabla^* p^* + \nabla^* \cdot \boldsymbol{T}^*, \\
\boldsymbol{T}^* &= \frac{1+\phi}{2}\boldsymbol{\sigma}_b^* + \frac{1-\phi}{2}\boldsymbol{\sigma}_w^* - \frac{\lambda}{g\mu H^2}\nabla^*\phi \otimes \nabla^*\phi, \\
\phi_{t^*}^* + \boldsymbol{v}^* \cdot \nabla^*\phi &= \frac{g(1-\phi^2)}{2} + \frac{\gamma_1 \lambda}{g H^4}\nabla^{*2}\left[-\nabla^{*2}\phi + \frac{\phi(\phi^2-1)}{(\epsilon/H)^2}\right], \\
\boldsymbol{\sigma}_{ext}^* &= 2\frac{\mu_{ext}}{\mu}\boldsymbol{D}^*, \\
\boldsymbol{\sigma}_b^* &= 2\boldsymbol{D}^* - (d-1)\boldsymbol{Q},
\end{aligned} \tag{5}$$

where we can identify a number of dimensionless groups. By relating the parameter $\lambda$, to an effective surface tension, $\sigma_{eff} = \frac{2\sqrt{2}}{3}\lambda/\epsilon$, we can rewrite some of the dimensionless groups in terms of a Capillary number, $Ca = \mu H g/\sigma_{eff}$. The resulting dimensionless groups and dimensional parameters are summarized in Table III.

At the start of the continuum simulations, the biofilm is a spherical cap with radius $r_0$ and origin $(0, 0, z_0)$. We took $z_0 < 0$, so that the initial shape was flatter than a hemisphere, to slightly better match the biofilm shape in experiments without breaking the initial symmetry in the $xy$-plane, although this did not affect the results strongly. The initial dimensional radius of the biofilm was assumed to be approximately 6 $\mu$m, in agreement with experimental measurements of biofilms at 10 h (see Fig. 3b in the main paper).

We solve the continuum equations (5) using the Dedalus Project [10]. Dedalus is a highly parallelized framework for solving partial differential equations. It uses pseudospectral methods for spatial discretization. We use Chebyshev polynomials in the wall-normal direction to enforce boundary conditions and Fourier polynomials in the streamwise and spanwise direction. The equations are integrated forwards in time using an implicit-explicit Runga-Kutta scheme. All models are run with resolution 384x196$^2$ in the streamwise, spanwise, and wall-normal directions.

| Parameter | Value | Unit | Description |
|---|---|---|---|
| $H$ | 40 | µm | Height of the channel. |
| $g$ | 1/8630 | $s^{-1}$ | Growth rate measured from experiments. For a volume $V$, a biofilm obeys $\dot{V} = gV$. |
| $Ca$ | 100 | | Effective capillary number. |
| $\epsilon/H$ | 0.01 | | Ratio of interface thickness and height of the channel. |
| $\mu_{ext}/\mu$ | 0.1 | | Ratio of biofilm and external viscosity. |
| $\gamma_1\mu/H^2$ | $10^{-4}$ | | Effective diffusivity of the phase-field. |
| $r_0/H$ | 0.17 | | Ratio of radius of initial spherical cap and height of the channel. |
| $z_0/H$ | -0.04 | | Ratio of the origin of initial spherical cap and height of the channel. |
| $\beta$ | $1.25 \cdot \frac{\pi}{2}$ | | Angle through which the director field rotates in the x-z plane. |
| $a_0/H$ | 0.325 | | Length of region over which the director field rotates at the back of the biofilm. |
| $a_1/H$ | 0.21 | | Length of region over which the director field rotates at the sides of the biofilm. |
| $e_sH/\mu$ | 0.5 | | Dimensionless slip parameter. |
| $S$ | 0.4 | | Strength of nematic order measured from experiments (see Fig. S7). |

TABLE III. Continuum simulation parameters.

#### EXPERIMENTAL DETAILS

Experiments were performed using *V. cholerae* strain KDV613, also referred to as WT*, which is a derivative of the N16961 wild type strain (O1 El Tor). Although *V. cholerae* typically displays a slightly curved cell body, the strain KDV613 displays a straight cell shape, owing to the deletion of the *crvA* gene (VCA1075). This strain also carries the low copy number plasmid pNUT542, which harbors the gene coding for the superfolder green fluorescent protein (*sf-gfp*) under the control of a constitutively expressed synthetic promoter ($P_{tac}$). The constitutive fluorescent signal was used to segment the *V. cholerae* cells in biofilm images as described previously [1], using, the biofilm image analysis software BiofilmQ [11]. Further experimental results to complement those in the main paper are shown in Figs. S10 and S11.

##### Calculations for quantitative definitions of phases

Quantitative definitions of each phase are shown in Fig. 1a. The volume of the shell is the flow-facing surface area of the convex hull multiplied by the cell width. Here the flow-facing surface area is the difference between the total surface area and the surface area of the base (calculated by taking the 2D convex hull in the $xy$ plane). The volume of the core is the remaining volume of the convex hull or zero, whichever is greater. For these calculations, the 10% of cells furthest from the biofilm center of mass were removed to reduce noise and artifacts.


[1] R. Hartmann, P. K. Singh, P. Pearce, R. Mok, B. Song, F. Díaz-Pascual, J. Dunkel, and K. Drescher, Nat. Phys. **15**, 251 (2019).
[2] T. Price, Q. J. Mech. Appl. Math. **38**, 93 (1985).
[3] G. B. Jeffery, Proc. Royal Soc. A **102**, 161 (1922).

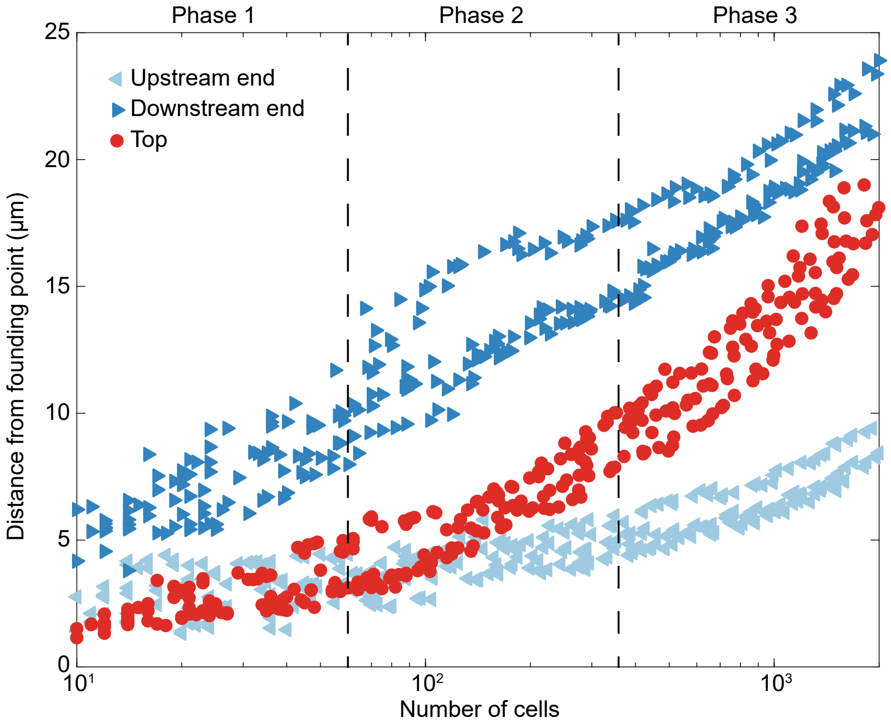


FIG. S8. Expansion characteristics in each growth phase. Each marker shows a distance from the founding point versus number of cells at a single timepoint ($n = 3$ biofilms). The distances were calculated by taking the minimum or maximum cell position in each direction, after removing the 10% of cells furthest from the biofilm center of mass, to reduce noise and artifacts. In phase 1, the biofilm expands fastest downstream. In phase 2, it begins to expand faster vertically, as cells begin to verticalize. In phase 3, most of the cells are vertical and it expands fastest upwards, and roughly equally fast in the upstream and downstream directions.


[4] T. Zhang, N. G. Cogan, and Q. Wang, SIAM J. Appl. Math. **69**, 641 (2008).

[5] K. Drescher, J. Dunkel, C. D. Nadell, S. van Teeffelen, I. Grnja, N. S. Wingreen, H. A. Stone, and B. L. Bassler, Proc. Natl. Acad. Sci. U.S.A. **113**, E2066 (2016).

[6] P.-G. De Gennes and J. Prost, *The Physics Of Liquid Crystals (International Series Of Monographs On Physics)* (Oxford University Press, 1995).

[7] A. Doostmohammadi, J. Ignés-Mullol, J. M. Yeomans, and F. Sagués, Nat. Commun. **9**, 3246 (2018).

[8] R. B. Bird, C. F. Curtiss, R. C. Armstrong, and O. Hassager, *Dynamics of polymeric liquids: Kinetic theory* (Wiley, 1987).

[9] P. Yue, J. J. Feng, C. Liu, and J. Shen, J. Fluid Mech. **515**, 293 (2005).

[10] K. J. Burns, G. M. Vasil, J. S. Oishi, D. Lecoanet, and B. P. Brown, arXiv:1905.10388 (2019).

[11] R. Hartmann, H. Jeckel, E. Jelli, P. K. Singh, S. Vaidya, M. Bayer, L. Vidakovic, F. Diaz-Pascual, J. C. Fong, A. Dragos, O. Besharova, C. D. Nadell, V. Sourjik, A. T. Kovacs, F. H. Yildiz, and K. Drescher, bioRxiv:735423 (2019).

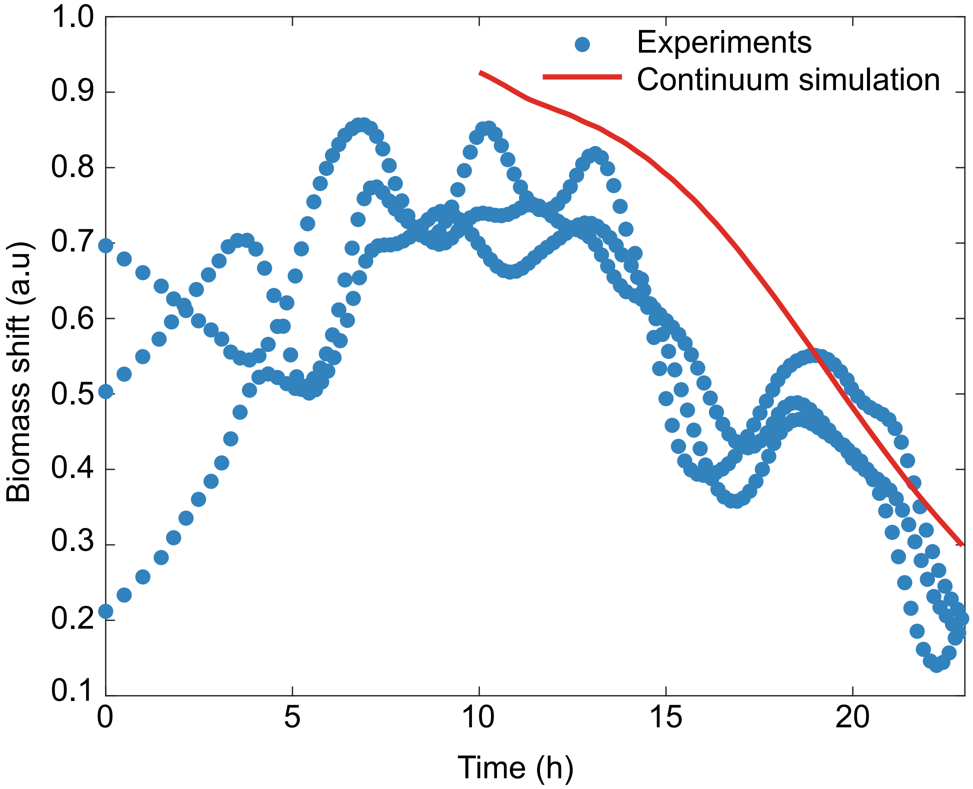


FIG. S9. Biomass shift versus time from experimental measurements ($n = 3$ biofilms) and continuum simulations. Biomass shift is defined as the sum of the biomass flux (shown in Fig. 3b of the main paper) along the flow direction, normalized by the sum of the absolute values, as described in [1].

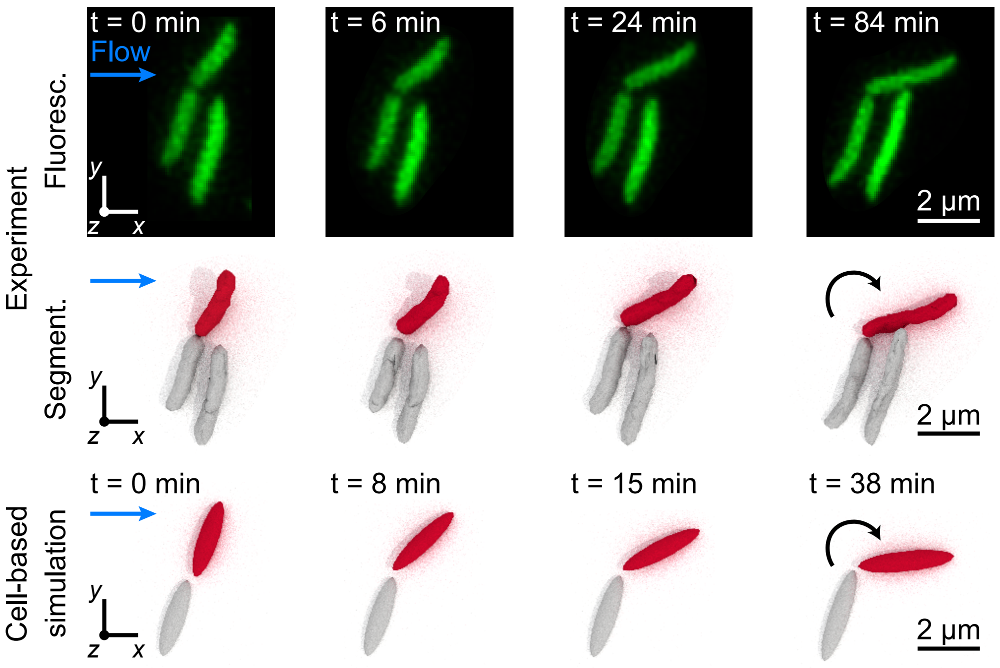


FIG. S10. Intermediate time steps in rotation of cell shown in Fig. 1b.

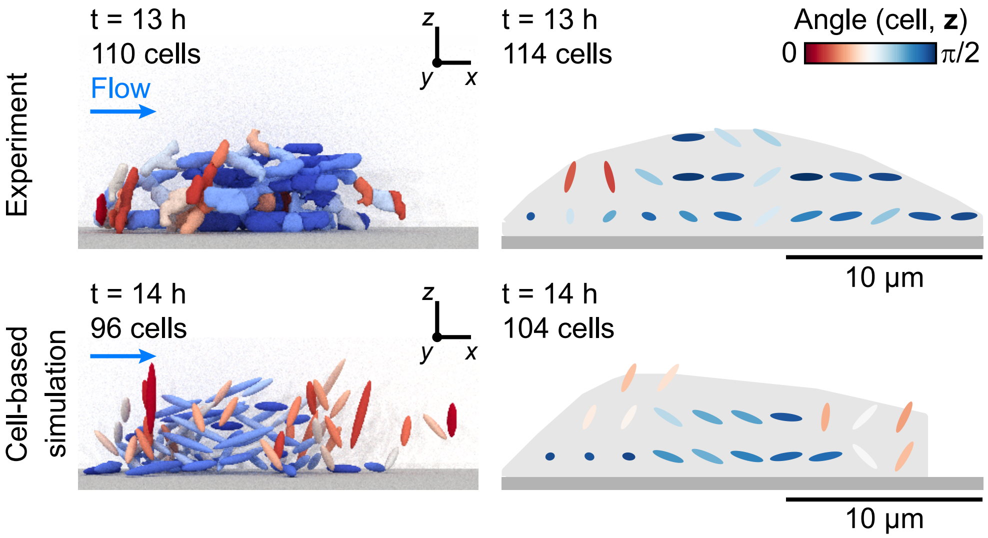


FIG. S11. In the early stages of growth, cells are predominantly aligned with the flow, with some vertical cells on the upstream side of the biofilm. The right-hand panels show averaged nematic alignment fields for several experiments ($n = 3$, top) and simulations ($n = 10$, bottom) along the midplane of the biofilm. In each case, the grey area denotes the region inside the convex hull around gridpoints with a cell number density higher than 0.05 $\mu m^{-3}$ per biofilm. The cell-based simulations recreated the cell alignment fields, consisting of mainly flow-aligned cells with some vertical cells at the front of the biofilm, suggesting that an applied shear is sufficient to explain their observation in our experiments.
